# DYRKP kinase regulates cell wall degradation in Chlamydomonas by inducing matrix metalloproteinase expression

**DOI:** 10.1101/2024.02.09.579704

**Authors:** Minjae Kim, Gabriel Lemes Jorge, Moritz Aschern, Stéphan Cuiné, Marie Bertrand, Malika Mekhalfi, Jae-Seong Yang, Jay J. Thelen, Fred Beisson, Gilles Peltier, Yonghua Li-Beisson

## Abstract

The cell wall of plants and algae is an important cell structure that protects cells from changes in the external physical and chemical environment. This extracellular matrix composed of polysaccharides and glycoproteins, is needed to be remodeled continuously throughout the life cycle. However, compared to matrix polysaccharides, little is known about the mechanisms regulating the formation and degradation of matrix glycoproteins. We report here that a plant kinase belonging to the dual-specificity tyrosine phosphorylation-regulated kinase (DYRK) family present in all eukaryotes regulates cell wall degradation in the model microalga *Chlamydomonas reinhardtii* by inducing the expression of matrix metalloproteinases (MMPs). In the absence of DYRKP, daughter cells fail to degrade the parental cell wall, and form multicellular structures. On the other hand, the complementation line of DYRKP was shown to degrade the parental cell wall normally. Transcriptomic and proteomic analyses indicate a marked down-regulation of MMP expression in the *dyrkp* mutants. Additionally, the expression of MMP was confirmed to be consistent with the expression pattern of DYRKP. Our findings show that DYRKP, by ensuring timely MMP expression, enables the successful execution of the cell cycle. Altogether, this study provides new insight into the life cycle regulation in plants and algae.

**IN A NUTSHELL:** *Background:* Plants and algae have different types of polysaccharides in their cell walls, but they have glycoproteins in common. Glycoprotein synthesis and degradation must be tightly regulated to ensure normal growth and differentiation. However, little is known about the regulatory mechanism of glycoprotein degradation in both plants and algae. The cell cycle of *Chlamydomonas reinhardtii* begins anew with the hatching of daughter cells, and the role of matrix metalloproteinases (MMPs) is known to be important in this process. In our previous study, we observed that a knockout mutant of the plant kinase belonging to the dual-specificity tyrosine phosphorylation-regulated kinase (DYRKP) formed a palmelloid structure and failed to hatch.

*Questions:* What is the role of DYRKP in microalgae? Specifically, why does the *dyrkp* mutant form a palmelloid structure? Palmelloid is usually observed in dividing cells or after exposure to stresses. We therefore hypothesized that the palmelloid phenotype observed in *dyrkp* mutant could either be due to a defect in cell hatching or due to an increased stress state in the mutant population.

*Findings:* We answered these questions by comparative studies in different culture conditions and by examining additional *dyrkp* knockout mutants generated by CRISPR-Cas9 in various background strains with more or less intact cell walls. Palmelloid formation in the *dyrkp* mutant was observed under optimal growth (mixo- or auto-trophic condition) and very low light conditions. Interestingly, unlike the parent strain, in which only cell wall fragments are observed in old cultures, the parental cell wall of the *dyrkp* mutant remained almost intact even after the release of daughter cells. Also, the cell division rate of the cell wall-less *dyrkp* mutants was similar to their background strain. These results suggest that *dyrkp* mutants have a problem in degrading the parental cell walls. Indeed, proteomic and transcriptomic analyses revealed reduced levels of protease families in the *dyrkp* mutant, and in particular with a significantly lower amount of several key members of the MMP family. Through the analysis of complementation lines, we confirmed that the DYRKP was required for strong and rapid expression of MMPs.

*Next steps:* We are pursuing research to understand what the phosphorylation clients of DYRKP are and how they regulate the expression of the MMPs identified in this study.

*One sentence summary:* The DYRKP kinase induces the expression of matrix metalloproteinases involved in the degradation of the parental cell wall, allowing prompt hatching of daughter cells after cell division.

## Introduction

The cell wall, one of the extracellular matrix (ECM) in plants and algae (Cosgrove, 2005; Domozych and LoRicco, 2023), is a complex network of molecules surrounding cells and tissues, providing them with mechanical support and transmitting regulatory cues from the environment (Gu and Rasmussen, 2022). The ECM, composed of matrix polysaccharides and matrix glycoproteins, is continuously synthesized and degraded during cell growth, division, and differentiation (Flinn, 2008; Seifert and Blaukopf, 2010; Domozych and LoRicco, 2023). Many studies about the regulatory mechanism for cell wall remodeling focus on matrix polysaccharides (Bashline et al., 2014; Anderson and Kieber, 2020), but the regulation for the synthesis and degradation of matrix glycoproteins has been rarely described.

In both animal and plant cells, glycoprotein hydrolyzing enzymes involved in ECM remodeling, called ECM proteases, are known to be essential for normal growth and development (Holmbeck et al., 1999; Golldack et al., 2002; Bonnans et al., 2014; Mishra et al., 2021); the loss of function of ECM proteases results in growth arrest (Holmbeck et al., 1999; Golldack et al., 2002), while overexpression of ECM proteases accelerates cancer invasion in animal cells (Tryggvason et al., 1987) or leaf senescence in plant cells (Wu et al., 2022). The control of ECM protease expression by kinase cascade (e.g., mitogen-activated protein kinase (MAPK)) has been well documented in animal cells (Wagner and Nebreda, 2009; Kumar et al., 2010), but only limited evidence is available for plant cells; MAPK3 and MAPK6 cascade induce the expression of matrix metalloproteinases (MMPs) during senescence in leaves (Wu et al., 2022).

*Chlamydomonas reinhardtii* (hereafter Chlamydomonas) is a photosynthetic unicellular microalga that has a non-cellulosic cell wall composed of hydroxyproline-rich glycoproteins, arabinose, mannose, and galactose (Baudelet et al., 2017; Goodenough and Lee, 2023). The formation and degradation of cell walls occur actively during Chlamydomonas cell hatching when daughter cells are released from the parental cell wall after mitosis (Cross and Umen, 2015). The MMPs (named gamete lytic enzymes) and the subtilisin-like serine proteases (SUBs; named vegetative lytic enzymes) are reported to be involved in cell wall degradation in Chlamydomonas (Kubo et al., 2009; Zou and Bozhkov, 2021). However, how cell wall degradation and ECM proteases are regulated in this model organism which displays both plant and ancestral eukaryotic features remains largely unknown.

In this study, we present the plant kinase belonging to the dual-specificity tyrosine phosphorylation-regulated kinase (DYRK) as a crucial regulator for algal cell wall degradation. Our comprehensive analysis demonstrates this kinase enables the successful execution of the cell cycle by ensuring timely *MMP* expression.

## Results

### The *dyrkp* mutant forms palmelloids independently of stress

In a previous study on the Chlamydomonas mutant *starch degradation 1 (std1,* called *dyrkp* mutant hereafter) which accumulates starch under nitrogen starvation conditions, we showed that this mutant has an insertional mutation in the plant DYRK (DYRKP) (Schulz-Raffelt et al., 2016). The DYRKP belongs to the DYRK kinase family which is known to be regulators of cell growth and development in eukaryotes (Laguna et al., 2008; Becker, 2012; Kurabayashi and Sanada, 2013). DYRK kinases are activated by autophosphorylation of conserved tyrosine residues and then phosphorylates serine/threonine residues of substrates (Aranda et al., 2011). In *dyrkp* mutant, we observed that cells tended to form clusters in liquid culture (**Fig. 1A**) (Schulz-Raffelt et al., 2016). Investigating this aggregation phenotype further, we found that the average particle size of the *dyrkp* mutant was 2**–**3 times bigger than its background strain 137AH (**Fig. 1B**). Within the *dyrkp* mutant population, multiple cells were trapped in a big cell wall (**Fig. 1C**), a structure usually called a palmelloid in literature (Goodenough and Lee, 2023). Cell population analysis revealed that the 137AH strain consisted of single cell mostly, whereas, more than 80% of the *dyrkp* mutant population consisted of multiple cells (**Fig. 1D**).

**Figure 1.**
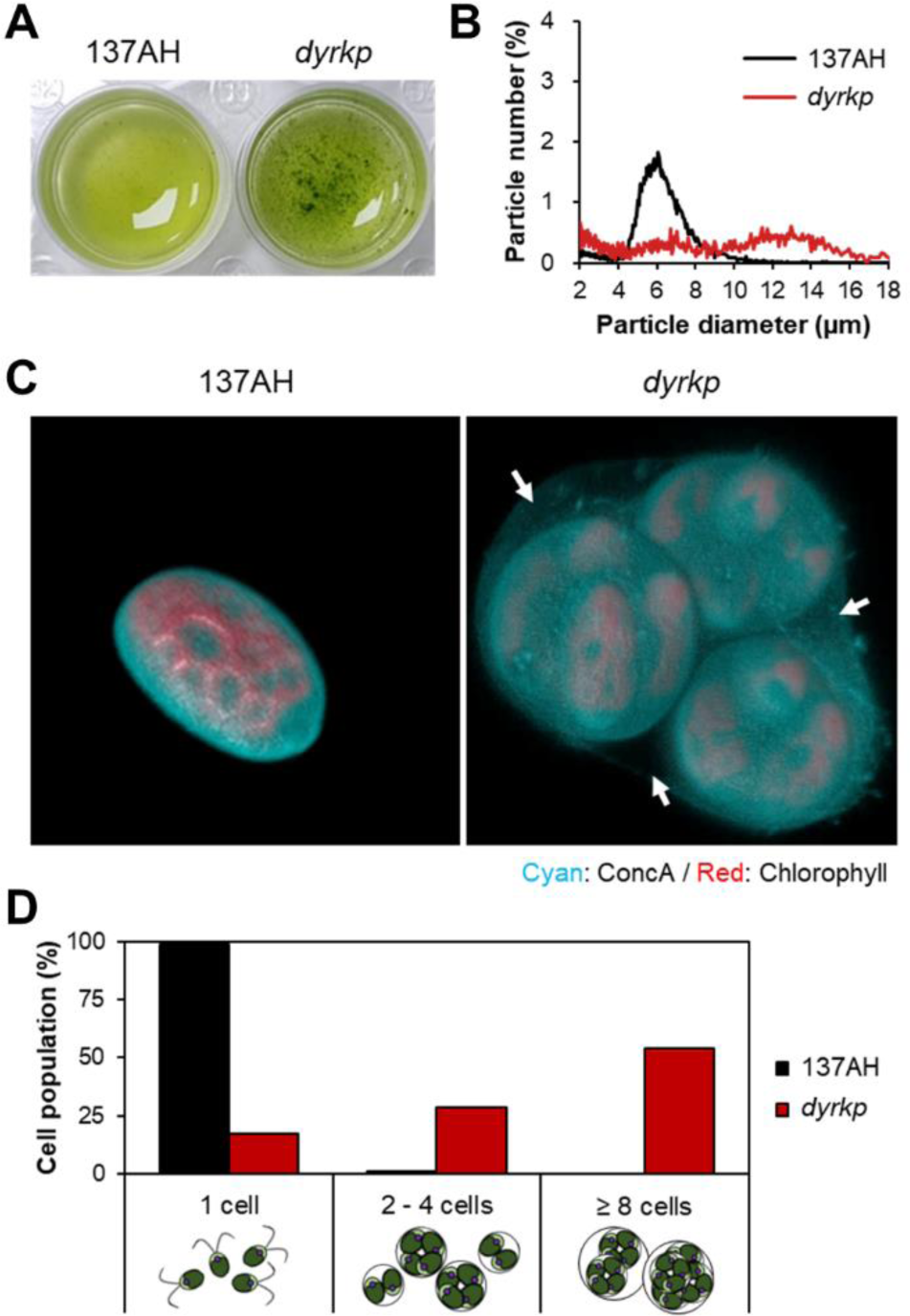
The *Chlamydomonas reinhardtii dyrkp* mutant formed palmelloid structure. **A)** Cell culture images. **B)** Distribution of particle size in the culture 3 days after inoculation. The mean value of particle size was calculated from all particles. **C)** Confocal microscopic images. The merged images indicate the cell wall stained with Concanavalin A conjugating fluorescent dye at 594 nm (ConcA; cyan) and the chloroplast with chlorophyll autofluorescence at 488 nm (Chl; red). The Z-stack images clearly showed that the *dyrkp* mutant cells were trapped inside three parental cell walls surrounded by another big cell wall (white arrows). **D)** Cell population distributions. The composition of particles classified into a single cell and the number of cells inside the parental cell wall. The sample was measured in the culture 3 days after inoculation.

During the asexual cell cycle, the particle diameter increases as palmelloid formation during mitosis, and as daughter cells hatch, the particle diameter decreases and particle concentration increases (**Supplemental Fig. S1A**). To measure growth, both 137AH strain and *dyrkp* mutant were treated with autolysin to release single cells and inoculated with the same particle number. Due to palmelloid formation, the growth pattern was expressed in particle concentration indicating the number of particles in volume, including single and multiple cells. The particle concentration of the *dyrkp* mutant increased slower than 137AH strain and became similar 14 d after inoculation under standard light condition (**Supplemental Fig. S1B**). Consistent with the slower increase in particle concentration, the decrease in mean particle diameter in *dyrkp* mutant was slower than in the 137AH strain (**Supplemental Fig. S1B**).

Chlamydomonas cells form palmelloids not only during mitosis but also when the cells are exposed to stress conditions such as high light (**Supplemental Fig. S1A**) (Khona et al., 2016; de Carpentier et al., 2019). To rule out that high light stress could be a trigger for the formation of palmelloids in *dyrkp*, we investigated cell morphologies under standard and very low light conditions. The particle size of the *dyrkp* mutant and 137AH strain increased in a similar manner under very low light condition (**Supplemental Fig. S1C**), suggesting that the palmelloid structure observed in the *dyrkp* mutant is not the consequence of stress responses. Since the palmelloids were observed in the *dyrkp* mutant regardless of nutritional conditions (**Supplemental Fig. S1D**), all subsequent studies were conducted under photoautotrophic conditions.

### Degradation of the parental cell wall is compromised in the *dyrkp* mutant

The total volume of the *dyrkp* mutant particles was up to three times bigger than that of 137AH (**Fig. 2A**). This phenotypical difference was also illustrated by the pellet size after low-speed centrifugation, which was much larger in the *dyrkp* mutant than in 137AH (**Fig. 2B**). Many undigested parental cell walls were observed in the *dyrkp* mutant (**Fig. 2C**). The undigested parental cell wall of the *dyrkp* mutant was clearly distinguished from the 137AH strain. From the ‘day 25’ culture of the *dyrkp* mutant, we observed the occurrence of a population with a smaller size in the cell counter, and this population disappeared after autolysin treatment, indicating that this population was composed essentially of the undigested parental cell wall (**Fig. 2D**). Next, we quantified and compared the total proteins present in the upper phase of the culture. We observed that the protein concentration of the culture medium increased rapidly in strain 137AH throughout the growth, but not of the *dyrkp* mutant (**Fig. 2E**). These results indicate the absence of DYRKP resulted in a delayed degradation of the parental cell wall.

**Figure 2.**
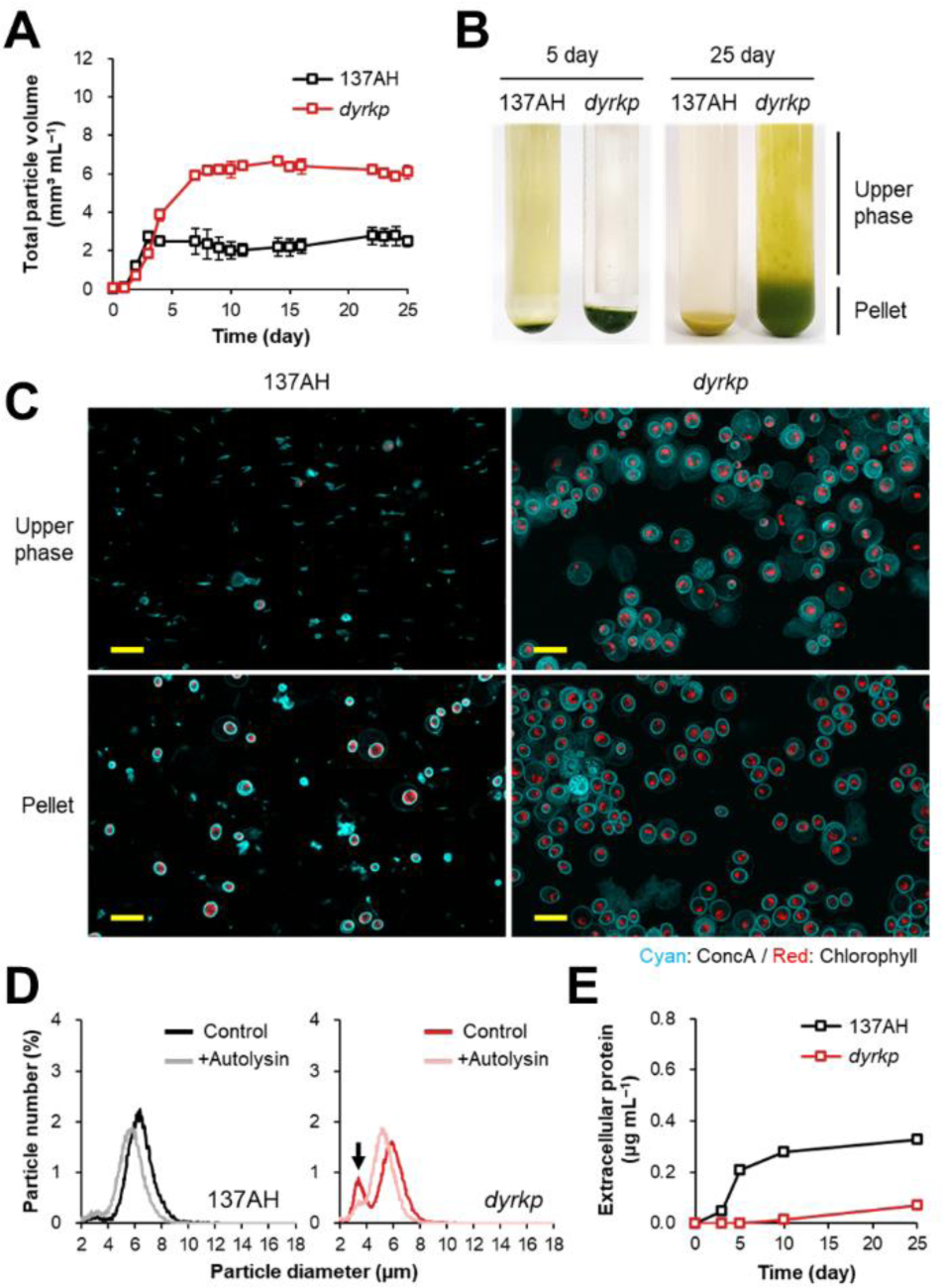
The *dyrkp* mutant showed an impairment in the digestion of the parental cell wall. **A)** Changes in the total particle volume under standard light condition (80 ± 5 μmol photons m^−2^ s^−1^). **B)** Morphology of pellets after low-speed (500 *g*) centrifugation. The same volume (10 mL) was collected from the culture of 5 days and 25 days after inoculation. **C)** Confocal microscopy images of the upper phase and pellet after low-speed centrifugation. The 25-day-old cultures were harvested and the samples were stained with the solution of concanavalin A conjugating fluorescent dye (ConcA; cyan) without filtration. Cells and empty parental cell walls can be distinguished by the presence of chlorophyll autofluorescence (Chl; red). The ConA and Chl signals were obtained at 594 nm and 488 nm, respectively. The yellow scale bar indicates 20 µm. **D)** Distribution of particle size in the 25-day-old cultures. The population was compared between the autolysin treatment (+Autolysin) and no treatment (Control) groups after 30 min of treatment. The black arrow points to the population of undigested parental cell wall debris. **E)** The amount of extracellular proteins in the culture medium. All experiments were performed at least in three biological replicates (± S.D).

We then investigated two complemented *dyrkp* mutant lines (DYRKP-c1 and DYRKP-c2; **Fig. 3A**) driven by the light-inducible *psaD* promoter (Schulz-Raffelt et al., 2016). The complemented lines formed intact and smaller size pellets than the *dyrkp* mutant (**Fig. 3B**) and their particle size decreased to a single cell size within seven days of continuous light conditions (**Fig. 3C**). However, unlike the 137AH strain, the particle concentration of complemented lines decreased rapidly after saturating point and the decline of the DYRKP-c2 line were faster than the DYRKP-c1 line (**Fig. 3D**). Based on these results, we hypothesized the DYRKP involves in the degradation of the parental cell wall.

**Figure 3.**
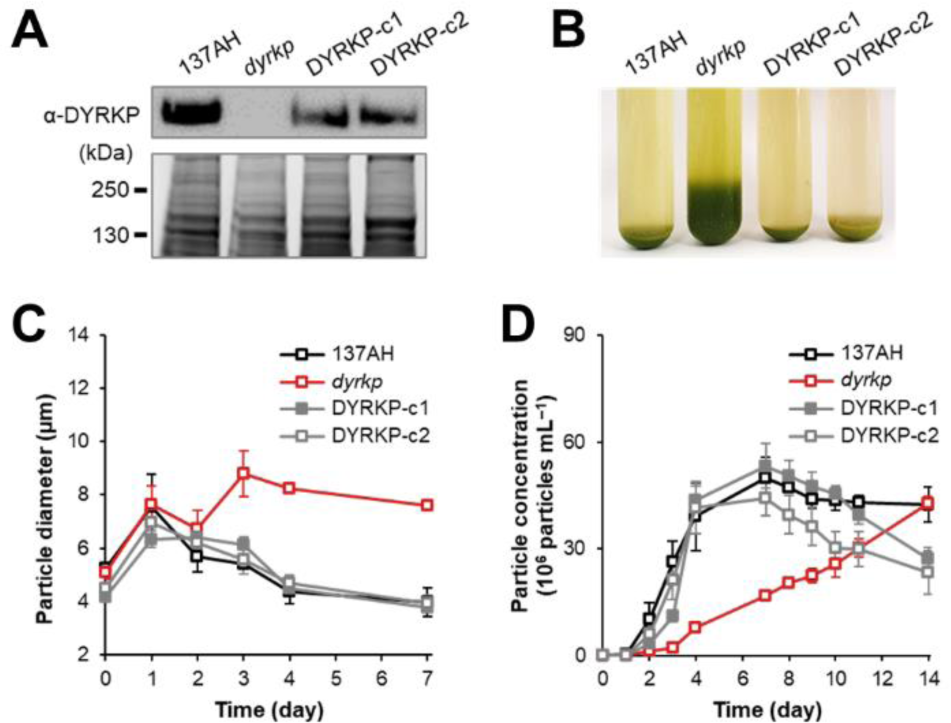
The complemented lines of *dyrkp* mutant (DYRKP-c1 and DYRKP-c2) appeared the restoration of parental cell wall digestion ability. **A)** Immunoblot of 137AH strain, *dyrkp* mutant, and two complemented lines. The upper and below panels indicated DYRKP antibody (α-DYRKP) and loading control, respectively. **B)** Morphology of pellets after low-speed (500 *g*) centrifugation. The same volume (10 mL) was collected from the culture of 25 days after inoculation. **C)** Changes in mean particle diameter under standard light condition. **D)** Changes in particle concentration under standard light condition; All experiments were performed at least in three biological replicates (± S.D).

### Palmelloid formation by the *dyrkp* mutant depends on cell wall integrity

137AH strain has a complete cell wall. To test our hypothesis, we sought to investigate the palmelloid phenotype of the *dyrkp* mutants derived from other cell-walled or cell-wall-less background strains. We obtained two mutants (*dyrkp2* and *dyrkp3*) of the CC5325 strain from the Chlamydomonas mutant library (Li et al., 2019). We generated target-specific DYRKP knockout mutants using the CRISPR-Cas9 method in the cell-walled strain CC125 (*dyrkp13*, *dyrkp15*, and *dyrkp24*) and in two cell-wall compromised strains CC4349 (*dyrkp3*, *dyrkp9*, and *dyrkp10*) and *dw15* (*dyrkp7* and *dyrkp10*). Since so-called cell wall defective strains have often various degrees of defects in their cell wall, we therefore first tested the cell wall integrity of the background strains using a detergent-treatment test (**Fig. 4A**). The insertion of the antibiotic cassette was confirmed by PCR analysis of the respective genomic DNA (**Supplemental Fig. S2**), and all mutants were confirmed to be true knockout by immunoblot analysis using anti-DYRKP (**Fig. 4B**).

**Figure 4.**
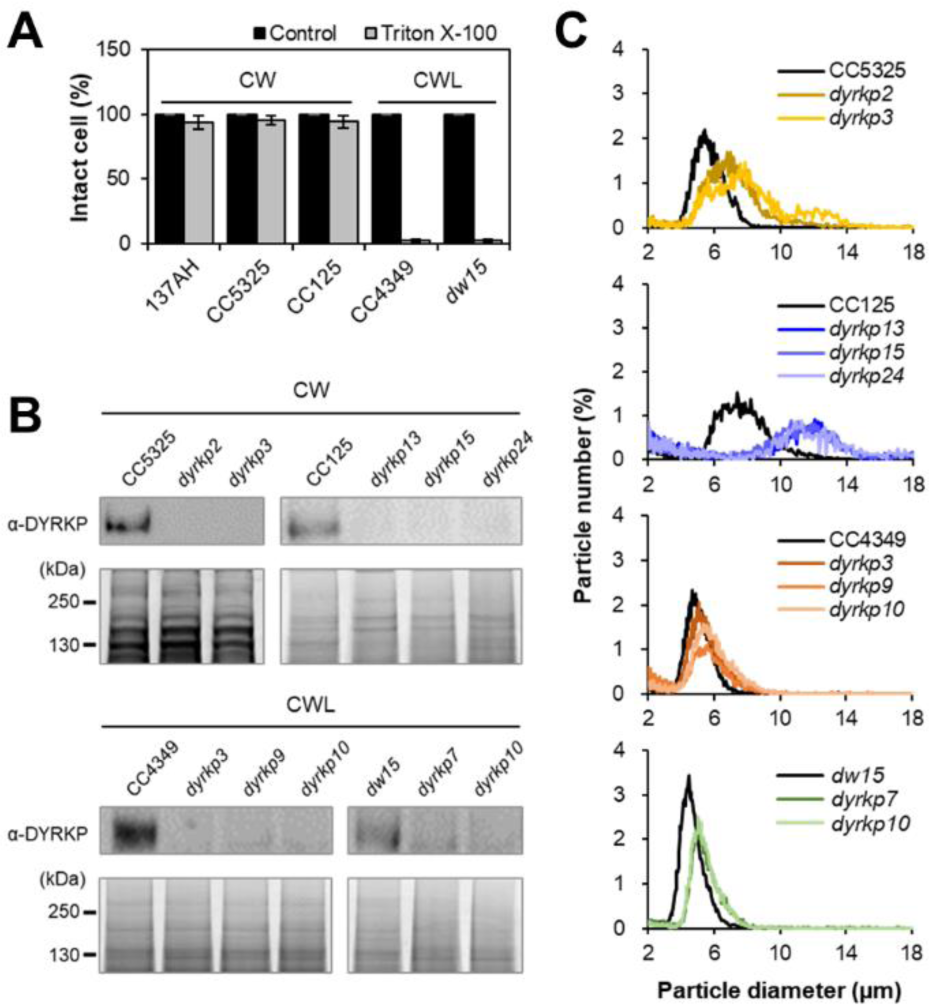
The *dyrkp* mutants generated from other cell-walled background strains formed palmelloid structures. **A)** Cell wall integrity was tested by detergent (Triton X-100) treatment in cell-walled (CW) and cell-wall-less (CWL) background strains. **B)** Immunoblot to DYRKP antibody (α-DYRKP) in the upper panel and loading control in the below panel. **C)** Distribution of particle size in the culture 3 days after inoculation.

Palmelloids were not observed in the *dyrkp* mutants derived from cell-wall-less strains (**Fig. 4C**; **Supplemental Fig. S3**), supporting our hypothesis that the mutant phenotype is related to the cell wall. Next, to understand the effect of DYRKP on the cell cycle, growth was measured under diurnal cycles (12 h light and 12 h dark). All *dyrkp* mutants showed a similar maximum cell density and maximum growth rate (**Supplemental Fig. S4**), indicating that DYRKP mutation does not affect the cell cycle.

On the other hand, the particle size of the *dyrkp* mutants in cell-walled strains was bigger than their background strains and the difference was more obvious in the CC125 background than in the CC5325 background (**Fig. 4C**). The CC5325-background mutants did not show large multicellular structures as much as those derived from 137AH or from CC125 (**Supplemental Fig. S3; Supplemental Fig. S5A**). The ratio of single cells in CC5325-background mutants was over 50%, while CC125-background mutants showed less than 25%, which was similar to the *dyrkp* mutant (**Supplemental Fig. S5B**). The changes in particle size of CC125-background mutants were more distinct than that of CC5325-background mutant (**Supplemental Fig. S5C-E**). Although the CC5325-background mutants formed an intact pellet unlike the *dyrkp* mutant and the CC125-background mutants (**Supplemental Fig. S5F**), a particle analysis showed a clearly different pattern to their background strains in common (**Supplemental Fig. S5G**). Overall, the palmelloid phenotype of the *dyrkp* mutant was greatly affected by the integrity of the cell wall, and the size of the palmelloid structure (i.e. the extent of the phenotype) was found to be associated to the degree of the defects in the cell wall of the background strains from which the mutant is derived.

### The *dyrkp* mutant secretes fewer cell wall proteins

To further pinpoint the role of DYRKP in cell wall degradation, we compared the protein profiles between 137AH and *dyrkp* mutant by separating extracellular proteins on SDS-PAGE followed by silver staining. The proteins collected from the upper phase of the ‘day 25’ culture revealed distinct protein patterns between *dyrkp* mutant and the 137AH strain (**Supplemental Fig. S6A**). To further understand the cause of the incomplete cell wall degradation of the *dyrkp* mutant, we carried out a global proteomic analysis of cellular (hereafter pellet) and extracellular proteins in the culture medium (hereafter upper phase) of 137AH and the *dyrkp* mutant (**Fig. 5A**, **Supplemental Fig. S6, B and C**). A total of 5,088 pellet proteins and 3,141 upper phase proteins were detected in the two strains, and after elimination of redundant proteins, the remaining 2,369 pellet proteins and 1,108 upper phase proteins were further analyzed (**Supplemental Fig. S6D, Supplemental Table S1 and S2**). The Venn diagram showed the unique and common proteins among the two strains (**Supplemental Fig. S6E**). Among protein groups with significant differences between 137AH and *dyrkp* mutant, 416 proteins in the pellet and 319 proteins in the upper phase were identified (**Fig. 5B**, **Supplemental Table S3 and S4**).

**Figure 5.**
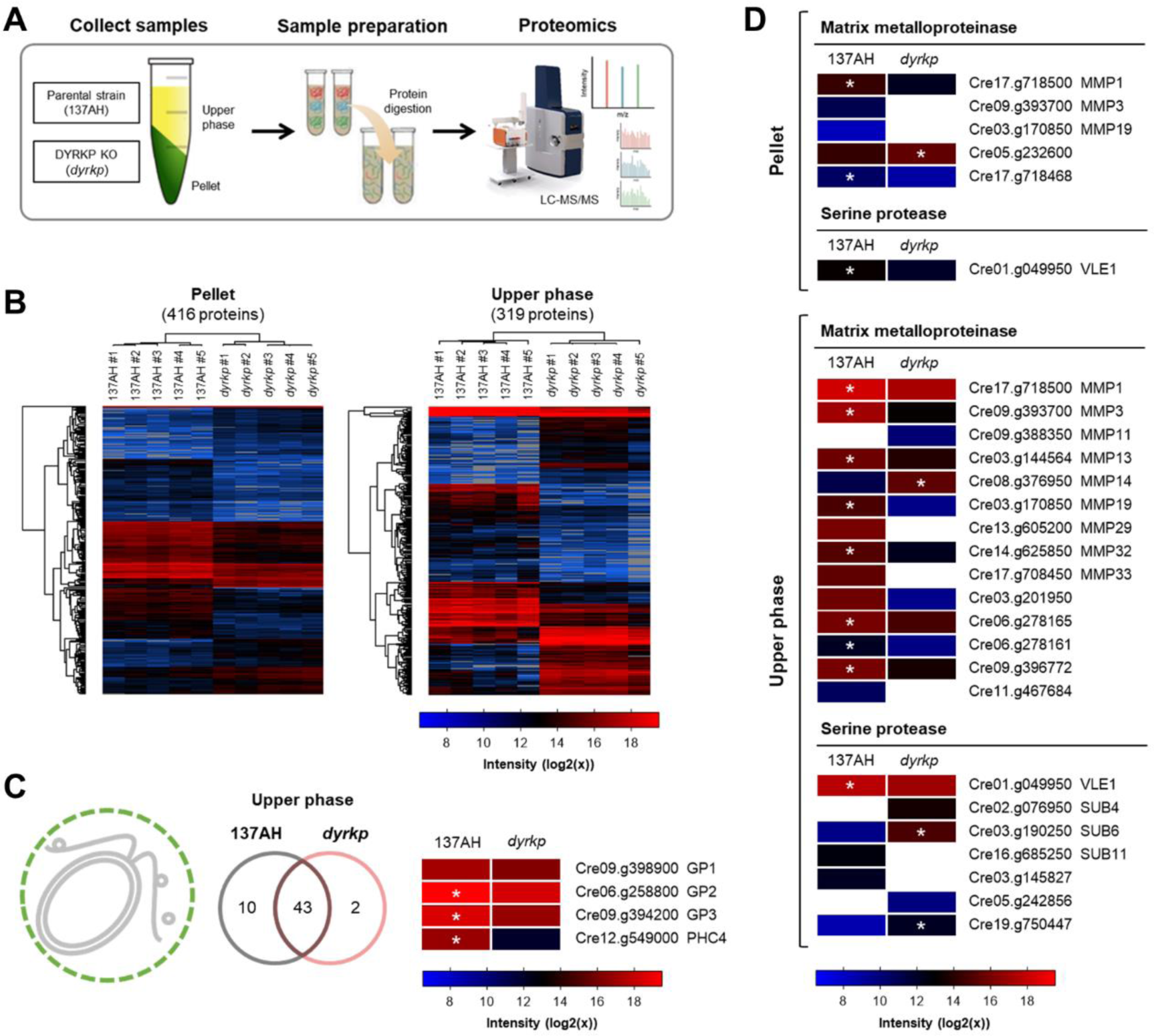
Cell wall proteins and ECM proteases were less abundant in the *dyrkp* mutant. **A)** Schematic diagram of the proteomic analysis workflow. The detailed analysis procedure is shown in **Supplemental Fig. S6**. **B)** Hierarchical clustering based on Euclidean distance. Five biological replicates in each group have high similarity to each other. **C)** The cell wall proteins detected in the upper phase. The cell wall proteins in the upper phase were assumed to be the cell wall proteins released from the parental cell walls. Venn diagram represents the number of commonly or uniquely found proteins between the 137AH strain and the *dyrkp* mutant. The heatmap shows the protein abundance of major cell wall proteins. Details are shown in **Supplemental Fig. S7**. **D)** Heatmap of the matrix metalloproteinases and serine proteases detected in the pellet and upper phase. Statistical analysis was performed using the non-parametric Mann-Whitney test; *p < 0.05 (± S.D).

In the extracellular medium (upper phase), 37 out of 43 cell wall-related proteins commonly detected in both strains were more abundant in the 137AH strain (**Fig. 5C**, **Supplemental Fig. S7**). This includes the well-known cell wall proteins of Chlamydomonas, i.e. glycoproteins (GP1, GP2, and GP3) and pherophorin-Chlamydomonas homolog 4 (PHC4) (Goodenough and Lee, 2023). These results therefore show that the low protein concentration in the culture medium of *dyrkp* mutant could explain the delay or defective degradation of the parental cell wall.

### Expression of ECM proteases is reduced in the *dyrkp* mutant

In Chlamydomonas, the parental cell wall is degraded by ECM proteases, which are transported through cilia (Long et al., 2016; Zou and Bozhkov, 2021) and secreted by ectosomes at the ciliary tips (Wood et al., 2013; Long et al., 2016). Cilia are essential for ectosome secretion because they are the only organelles not surrounded by a cell wall (Wood et al., 2013). Indeed, the delay or failure of cell hatching is observed in the cilia-assembly mutants of Chlamydomonas (Brown et al., 2015; Wang et al., 2022). Thus, we investigated the specific proteins associated with cilia and ectosome based on the previous proteome data (Long et al., 2016). Cilia-component proteins such as flagella-associated proteins (FAPs) and intraflagellar transport proteins (IFTs) were less abundant or even absent in the *dyrkp* mutant (**Supplemental Fig. S8A**). When the ciliary length was measured after autolysin treatment, it was observed that the *dyrkp* mutant had shorter cilia or even no cilia comparing to the 137AH strain (**Supplemental Fig. S8b**). Nevertheless, in the *dyrkp* mutant, a ciliary movement was observed from some daughter cells inside and outside of the parent cell wall (**Supplemental Fig. S8C, Supplemental Movie S1 to S5**), suggesting that lack of DYRKP delays cilia assembly but without affecting cell motility. For the ectosomal proteins, most of them were detected in both strains but were less abundant in the *dyrkp* mutant (**Supplemental Fig. S9**), indicating the delayed cilia assembly might have affected ectosome secretion.

The parental cell wall was still not degraded even after prolonged cultivation (**Fig. 2**), suggesting that there is likely a defect in the synthesis or transport of ECM proteases. Indeed, MMP1 and VLE1, major known ECM proteases of Chlamydomonas (Kubo et al., 2009; Zou and Bozhkov, 2021), were found in both strains but were more abundant in the 137AH strain than in the *dyrkp* mutant (**Fig. 5D**). Except for MMP14 and the protein product encoded by the locus Cre05.g232600, all other 13 MMP1 homologs were more abundant in the 137AH strain than *dyrkp*. On the other hand, of the six VLE1 homologs, only SUB11 and the protein product encoded by the locus Cre03.g145827 were enriched in the 137AH strain. Taken together, proteomic analysis indicates that the DYRKP affects cilia assembly as well as the amount of ECM proteases.

To identify difference in gene expression levels, transcriptomic analysis was performed. Compared to the 137AH strain, 687 up-regulated genes and 1568 down-regulated genes were identified in the *dyrkp* mutant (**Supplemental Fig. S10A, Supplemental Table S5**). Validation of transcriptome data by qPCR showed high accuracy (**Supplemental Fig. S10B**). Consistent with proteomics analysis, many hydrolytic enzymes for glycoproteins like MMPs and SUBs were down-regulated in the *dyrkp* mutant (**Fig. 6A**). Particularly, MMP and SUB showed a high positive correlation between transcriptome and proteome (**Supplemental Fig. S10C**). Among ECM proteases identified in the transcriptome and proteome, *MMP1*, *MMP3*, *MMP13*, *MMP19*, *VLE1*, and *SUB11* were selected and qRT-PCR further validated their differential expression pattern. As a result, the overall expression of the *MMP* family was significantly decreased in the *dyrkp* mutant but the expression of *VLE1* and *SUB11* did not show statistically significant differences (**Fig. 6B**). Similar patterns were confirmed in other cell-walled DYRKP knockout mutants (**Supplemental Fig. S11A**). The decreased expression of *MMP1*, *MMP3*, and *MMP13* were also observed in the cell-wall-less or -compromised DYRKP knockout mutants (**Supplemental Fig. S11B**).

**Figure 6.**
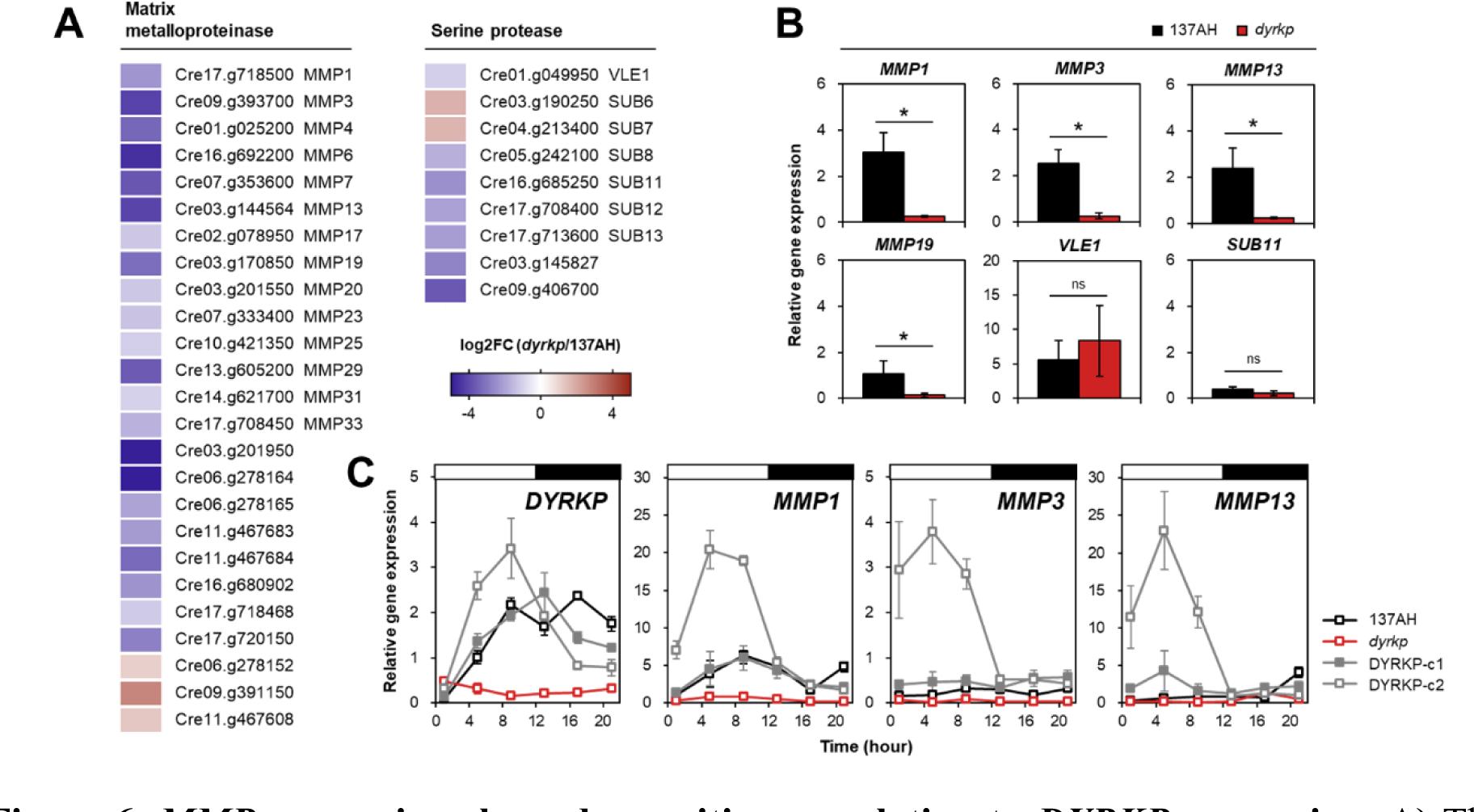
*MMP* expression showed a positive correlation to *DYRKP* expression. **A)** The differentially expressed genes encoding the matrix metalloproteinases and serine proteases. **B)** Gene expression levels of selected matrix metalloproteinases and serine proteases in 137AH strain and *dyrkp* mutant. **C)** Gene expression pattern of *DYRKP*, *MMP1*, *MMP3*, and *MMP13* in 137AH strain, *dyrkp* mutant, and two complemented lines (DYRKP-c1 and DYRKP-c2) under light/dark conditions. The value was normalized by the reference gene (*RACK1*). All experiments were performed at least in three biological replicates. Statistical analysis was performed using the non-parametric Mann-Whitney test; *p < 0.05 (± S.D). The ‘ns’ indicates no significance.

In addition to ECM proteases, down-regulated genes in the *dyrkp* mutant include those related to protein phosphorylation, kinase activity and phosphotransferase activity. Among 207 kinase genes, 177 were down-regulated in the *dyrkp* mutant. Among 22 phosphatase genes, 14 were down-regulated in the *dyrkp* mutant (**Supplemental Fig. S10D, Supplemental Table S5**).

### DYRKP positively regulates the expression of *MMP1*, *MMP3* and *MMP13*

Our multi-omics analysis indicated that DYRKP likely plays a role in regulating MMP expression. According to the gene expression database under light/dark conditions (Proost and Mutwil, 2018), *DYRKP* expression is associated with the life cycle of Chlamydomonas. In detail, the *DYRKP* expression increases gradually during the cell growth and peaks at the end of the light and dark phases which correspond to G1 and G0 phases (**Supplemental Fig. S12A**). Indeed, the expression in the 137AH strain increased once in each light and dark phase (**Fig. 6C**). However, DYRKP expression in the two complemented *dyrkp* mutants driven by the light-inducible promoter increased only in the light phase.

We then confirmed the expression patterns of *MMP1*, *MMP3*, *MMP13* and *VLE1* under light/dark conditions (**Fig. 6C**, **Supplemental Fig. S12B**). In the *dyrkp* mutant, the expression levels of *MMPs* were lower than the 137AH strain at almost all sampling points. In the 137AH strain, the expression of *MMP1* and *MMP3* increased during both light and dark phases while the expression of *MMP13* increased only in the dark phase. Similar to our results, gene expression databases indicated that the expression of *MMP1* and *MMP3* gradually increased and peaked at the end of each phase, and the expression of *MMP13* increased only in the dark phase. In the complemented lines, the expression of *MMP1*, *MMP3*, and *MMP13* was restored to a similar and even higher level than that in the 137AH strain, and their expression increased in the light phase and decreased in the dark phase, similar to the expression pattern of *psaD* gene. As expected, the expression of *VLE1* did not show a clear association with *DYRKP* expression.

## Discussion

Although the importance of cell hatching in the Chlamydomonas cell cycle is evident, little is known about its regulation. Here, we demonstrate that DYRKP plays an important role in cell hatching by regulating ECM degradation (**Fig. 7**). We further point out that this regulation is achieved through positively modulating the expression of ECM-associated MMPs. Genetic knockouts of DYRKP delayed the degradation of ECM components during cell division, and sustained expression of DYRKP in complemented lines restored the phenotype of the background strain. Moreover, many *MMPs* were repressed in the *DYRKP* knockout mutants, and the expression of *MMPs* altered consistent with the expression pattern of *DYRKP*.

**Figure 7.**
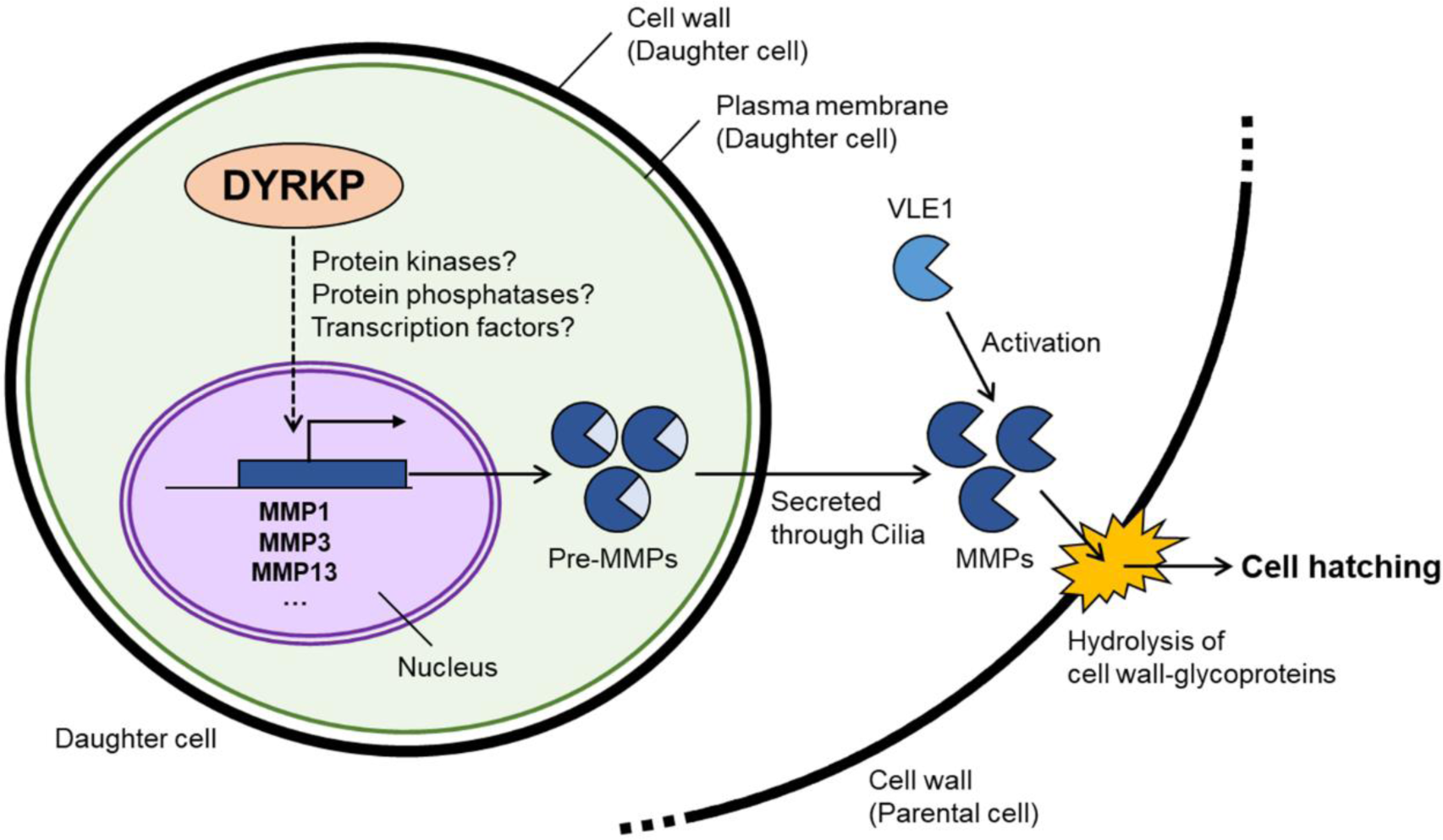
Schematic diagram of the role of DYRKP in cell hatching. DYRKP induces increased transcription of *MMP1*, *MMP3*, and *MMP13* genes located in nuclear DNA through phosphorylation of unknown protein kinases, protein phosphatases, or transcription factors. Afterward, the translated protein in the pre-activated form of MMPs (pre-MMPs) is delivered through cilia (Long et al., 2016; Zou and Bozhkov, 2021) and secreted in the form of an ectosome from the cilia membrane (Wood et al., 2013; Long et al., 2016). Pre-MMPs are activated by other proteases (Wilkinson et al., 2017); for example, pre-MMP1 is activated by VLE1 (Kubo et al., 2009; de Carpentier et al., 2022). Activated MMPs then hydrolyze and degrade the hydroxyproline-rich glycoprotein structure of the parental cell wall (Goodenough and Lee, 2023), causing fragmentation of the parental cell wall. When the degradation of the parental cell wall progresses sufficiently, the daughter cells are released (cell hatching), and the remaining parental cell wall is completely degraded by MMPs that remain active.

MMPs are important for a balanced and regulated degradation of ECM proteins during growth, morphogenesis and development in multicellular organisms (Holmbeck et al., 1999; Golldack et al., 2002; Page-McCaw et al., 2007). For example, the *At2-mmp* mutant of *Arabidopsis thaliana* is defective in tissue development, and its leaf cells are smaller than the wild-type and unevenly distributed (Golldack et al., 2002). Interestingly, the DYRKP knockout mutants of *M. polymorpha* failed to decompose the cilia-like structure usually degraded in wild-type strains (Furuya et al., 2021). In the case of Chlamydomonas, it has also been observed that the mutant defected in MMP32 exhibits a spontaneous aggregation phenotype (de Carpentier et al., 2022). The phenotypes of MMP-defective mutants are similar to those defected in DYRKP, further supporting our finding that DYRKP regulates cell hatching by controlling MMP expression.

Regulation of cell cycle progression by members of the DYRK family has been observed in animal cells at different stages of the cell cycle (Becker, 2012). For example, DYRK1A and DYRK1B regulate the length of the G1-phase as well as the decision between the entry and exit of the cell cycle; DYRK2 is a negative regulator of S-phase entry and is down-regulated in tumor cells; DYRK3 regulates the cell cycle of erythroblasts. On the other hand, DYRKP does not directly affect cell division (Furuya et al., 2021) but is involved in the last step of a cell cycle, i.e., cell hatching. In contrast to our study, DYRKs are found to act as a negative regulator of the cell cycle, highlighting the complexity and difference in the evolutionary trajectory of DYRKs in different kingdoms of life. Nevertheless, these observations collectively establish the conserved role of DYRKs in regulating the cell cycle in eukaryotes.

Expression of *MMP1*, *MMP3* and *MMP13* responded rapidly to DYRKP expression, thus it may be interesting to understand how this kinase regulates them simultaneously. In this regard, it is worth mentioning that many protein kinases and phosphatases were down-regulated in the *dyrkp* mutant. Among them, the aurora kinases which are the family of serine/threonine kinases, perform essential functions during cell division (Carmena and Earnshaw, 2003). In breast cancer cells, the siRNA-mediated silencing of aurora kinases markedly decreases the MMP-9 expression levels induced by the protein kinase C pathway (Noh et al., 2015). On the other hand, MAPKs induce the expression of MMPs during ECM remodeling and degradation in metastatic cancer cells (e.g. p38 MAPKs) (Wagner and Nebreda, 2009; Kumar et al., 2010) or senescence in leaves (e.g. MAPK3/MAPK6) (Wu et al., 2022). Another possibility is the phosphorylation of transcription factors. In animal cells, DYRK1 and DYRK2 are known to be involved in cancer initiation and progression such as proliferation and invasion by phosphorylating the nuclear factor of activated T cells (NFAT) (Shou et al., 2015). Direct regulation of MMP3 by NFAT1 promotes the growth and metastasis of melanoma tumors (Shoshan et al., 2016). However, the relationship between DYRK family and MMP in plants is still unknown, and it has not been confirmed whether DYRKP has the same phosphorylation client as DYRK1 and DYRK2. Therefore, further studies on direct DYRKP targets through phosphoproteomics or kinase client assays (Huang and Thelen, 2012) can help to understand its regulatory mechanism more clearly.

Taken together, our data broaden the context of DYRKP activity by demonstrating that this kinase has an impact on cell wall remodeling. Future work on candidates for DYRKP phosphorylation will provide a better understanding of how DYRKP is involved in regulating various cellular processes in plants and algae.

## Materials and Methods

### Strains and growth conditions

Background strains of Chlamydomonas were selected into two groups; cell-walled strains (137AH, CC5325, and CC125) and cell wall-less or cell wall-compromised strains (CC4349 and *dw15*). The DYRKP knockout mutant, named *std1* (*starch degradation 1*), was obtained from our mutant library (Schulz-Raffelt et al., 2016). All mutants and their background strains are shown in the **Supplemental Table S6**. Cells were cultured in 20 mL of photoautotrophic culture medium (MOPS-buffered minimal medium; MM-MOPS) in a 100 mL flask. The cells were cultured photoautotrophically in incubator shakers (Infos HT) with natural dissolving of 2% CO_2_ by continuous shaking (120 rpm) under continuous illumination (80 ± 5 μmol photons m^−2^ s^−1^) at 25 ± 1°C unless otherwise stated.

### Cell wall integrity assay

To confirm the cell wall integrity of background strains, 1 mL of 137AH, CC5325, CC125, CC4349, and *dw15* cultures was harvested by centrifugation at 1,000 *g*. The pellet was resuspended with 0.05% (w/v) Triton X-100 or with MM-MOPS medium by vortexed vigorously for 30 s then incubated in dark for 30 min. The size of the cell population and the number of intact cells were measured using a Coulter Counter (Multisizer 4; Beckman Counter, Brea, CA, USA).

### Generation of knockout mutants by CRISPR-Cas9 RNP method

To obtain *dyrkp* mutants, the CRISPR-Cas9 RNP method was used with a few modifications (Kim et al., 2020). The single guide RNA (sgRNA) sequence for the specific target sequence on the DYRKP gene was designed by Cas-Designer (http://www.rgenome.net/cas-designer) and selected as 5′-TCCTCGCAGCAAGTCTGCGC AGG -3′ (**Supplemental Fig. S2**). To increase the selection efficiency, insertion and expression of hygromycin-resistance (HygR; aphVII) gene cassette was combined. The purified Cas9 protein (100 μg, Cas9 expression plasmid: pET-NLS-Cas9-6xHis (Plasmid #62934)) and 70 μg of sgRNA synthesized by using GeneArt™ Precision gRNA Synthesis Kit (ThermoFisher, MA, USA), were mixed to form the sgRNA and Cas9 protein (RNP) complex. The RNP complex and aphVII gene expression cassette were co-transformed with electroporation (600 V, 50 μF, infinity resistance). After 12 hours of incubation to allow recovery from electroporation shock, cells were plated on a TAP medium containing 1.5% agar and hygromycin (25 μg mL^−1^; ThermoFisher). To screen *dyrkp* mutants, the colonies grown onto the selection plate were confirmed by genomic PCR using the target-specific primer sets (**Supplemental Table S7**).

### Microscopy analysis

For the light microscopic observation, the Leica DMRXA microscope (Leica Microsystems, Wetzlar, Germany) was used. To measure the ciliary length, cells were fixed with 0.2 % (w/v) glutaraldehyde added to the medium. Images and videos were obtained by using the Spot Insight 4 software (Diagnostic Instruments Inc., Sterling Heights, USA).

For the Confocal microscopic observation, the ZEISS LSM 780 Confocal microscope (Carl-ZEISS, Oberkochen, Germany) was used. To visualize cell wall structures, cells were stained with Concanavalin A (ConA) – Alexa Fluor^TM^ 594 conjugates (ThermoFisher) for 10 min at room temperature. The ConA was excited with a 561 nm laser and an emitting signal was collected between 582-635 nm. Chlorophyll autofluorescence was excited with a 488 nm laser and emission was collected between 648-683 nm. Data analysis was performed with the FIJI program. To quantify the cell population, all particles were classified into three groups; a single cell, 2-4 cells, and more than 8 cells. The range of chlorophyll autofluorescence signal area was determined based on more than 50 particles. To make sure our criteria of data analysis, the ratio was double-checked in the transmitted image. Afterward, all the particle signal from five independent images was classified into three groups and were expressed as a ratio.

### Quantitative and qualitative assays for extracellular proteins

Cell culture (2 mL) was collected from the culture after 0, 3, 5, 10, and 25 days of inoculation. To separate extracellular proteins and cellular proteins, cell cultures were centrifuged at 2,000 *g* for 5 min and the upper phase was carefully collected. Additionally, the collected culture medium was filtered with a 0.45 μm polyethersulfone (PES) membrane (VWR International, Radnor, PA, USA) to avoid contamination by the cellular proteins. Then, proteins in the filtered culture medium were precipitated by adding 5 volumes of ice-cold 100% acetone and were collected after centrifugation (10 min, 16,000 *g*, 4°C). After washing pellets three times with ice-cold 80% acetone, it was resuspended in 2 mL of 0.1% (w/v) SDS buffer and used for quantitative and qualitative assays.

For the quantitative assay, the total proteins in the samples were measured through the BiCinchoninic acid Assay using a kit (Sigma-Aldrich, Saint Louis, MO, USA). The assay was conducted according to the method provided by the manufacturer. For the qualitative assay, 100 ng of total proteins in ‘day 25’ were separated on a 10% (w/v) Bis-Tris gel (Invitrogen, Waltham, MA, USA) run for 1 h at 190 V in Novex Nupage (Invitrogen) and stained by the Silver staining method.

### Protein extraction and immunoblot analysis

Soluble cell extracts were prepared as follows: 2 mL of *C. reinhardtii* cell culture in the exponential phase (eq. to 5 × 10^6^ cells mL^−1^ or 0.8 mm^3^ mL^−1^) were harvested by centrifugation for 2 min at 1,789 *g* and resuspended in 0.5 mL lysis buffer (20 mM HEPES– KOH pH 7.2, 10 mM KCl, 1 mM MgCl_2_, 154 mM NaCl, 0.1× protease inhibitor cocktail). Cells were sonicated on ice for 30 s with an alternating cycle of 1 s pulse/1 s pause. After centrifugation (10,000 *g* 10 min 4°C), soluble proteins in the supernatant were precipitated with ice-cold 80% (v/v) acetone (−20°C, 1 h). Samples were then centrifuged for 10 min, 16,000 g at 4°C. The protein pellet was resuspended with Novex Nupage LDS buffer 1× (Invitrogen), and proteins were then denatured at 70 °C for 20 min. Protein extracts (30 µg protein) were loaded on 3-8% Novex Nupage Tris-Acetate gel (Invitrogen) and run for 1 h at 190 V in Novex Nupage (Invitrogen). Proteins were transferred onto nitrocellulose using a semidry transfer technique. Immunodetection of DYRKP was performed following the process described in our previous work (Schulz-Raffelt et al., 2016).

### Protein extraction and *in-solution* digestion

For proteomic analysis, the ‘day 5’ cultures of the 137AH strain and the *dyrkp* mutant were harvested by centrifugation (2,000 *g*) for 10 min. To ensure data reliability, samples were prepared from five biologically independent cultures. After centrifugation, the upper phase and pellet were carefully separated as the samples of extracellular proteins and cellular proteins, respectively. To avoid the contamination of cellular proteins and extracellular proteins, the isolated culture medium was filtered using a 0.45 μm PES membrane.

The pellet samples were extracted by adding 2.5 mL of Tris pH 8.8 buffered phenol and 2.5 mL of extraction media (0.1 M Tris-HCl pH 8.8, 10 mM EDTA, 0.4% (v/v) 2-mercaptoethanol, 0.9 M sucrose) and agitating for 30 min at 4°C. The phenol phase was removed, and the aqueous phase was back extracted with 2.5 mL of extraction media added with 2.5 mL of phenol. Phenol-extracted proteins were precipitated by adding 5 volumes of 0.1 M ammonium acetate in 100% methanol. On the other hand, the upper phase samples were precipitated right away without the phenol extraction process.

Precipitated proteins were collected after centrifugation (10 min, 16,000 *g*, 4°C). Pellets were washed twice with 0.1 M ammonium acetate in methanol, and three times with ice-cold 80% acetone. An aliquot was removed and centrifugated at 16,000 *g*, 4°C for 5 min followed by pellet resuspension in urea buffer (6M urea, 2M thiourea, 100 mM ammonium bicarbonate). Protein content was quantified by Bradford assay with Bovine Gamma Globulin as standard.

Before mass spectrometry analysis, 10 μg of each sample was aliquoted and normalized to the same concentration and volume (40 µL) in urea buffer (6M urea, 2M thiourea, 100mM ammonium bicarbonate). Reduction (10mM DTT in 10 mM ammonium bicarbonate) was performed at 30°C for 30 min and alkylation (40 mM iodoacetamide in 10mM ammonium bicarbonate) for 1 hour at room temperature in the dark. Samples were diluted to a final urea concentration of 0.8 M before tryptic digestion. Protein digestion was performed by adding a volume of trypsin aiming at an enzyme/protein ratio of 1:50 at 37 °C for 16 h. To achieve optimal digestion, a second addition of trypsin was done, and samples were incubated for 4 h. After digestion, the tryptic peptides were dried under vacuum centrifugation. Subsequently, samples were loaded onto Evotips according to the manufacturer’s instructions.

### Mass spectrometry data acquisition (UHPLC-MS/MS)

The acquisition of mass spectrometric data after tryptic digestion was performed on an ultra-high-performance liquid chromatography (UHPLC) EvoSep system that was used for reverse-phase liquid chromatography over a 44 min gradient at 250 nL min^-1^ flow. Peptides were separated on a C18 analytical column (PepSep C18 Bruker Daltonics, 15cmx150µm, 1.5µm particle size). Acquisition of mass spectra was carried out concomitantly to the chromatographic separation and peptides were analyzed through a Data Dependent Acquisition (DDA) method using a TimsTOF Pro 2 (Bruker Daltonics, Billerica, MA, USA). The instrument was operated in positive-ion, data-dependent PASEF mode over a m/z range of 100 to 1700. PASEF and TIMS were set to “on” for PASEF and TIMS. One MS and ten PASEF frames were acquired per cycle of 1.17sec. Target MS intensity was set at 10,000 with a minimum threshold of 2500 and from 20 to 59 eV, and a charge-state-based rolling collision energy table was employed. An active exclusion/reconsider precursor method with release after 0.4min was used. A second MS/MS spectra were obtained if the precursor had a four-time increase in signal strength in subsequent scans (within a mass width error of 0.015 m/z).

### Protein identification and data analysis for global proteomics

The PEAKS Studio 10.0 (Bioinformatics Solutions, Waterloo, Ontario, Canada) software was used to provide automated *de novo* sequencing from MS/MS spectra. PEAKS *de novo* sequencing was performed with precursor and fragment error tolerance values of 20 ppm and 0.01 Da, respectively. Trypsin was used as a protease with 3 maximum missed cleavage allowed. Carbamidomethylation of cysteine and oxidation of methionine were set as fixed and variable modifications, respectively. A maximum of four variable modifications per peptide was allowed. PEAKS DB, which is a database search module in PEAKS Studio, was used in the second step to identify peptide spectrum matches (PSMs) from existing protein databases (Zhang et al., 2012). The *C. reinhardtii* (19,526 entries) was used as the reference database. The target-decoy method known as “decoy fusion” that is included in PEAKS was utilized to estimate the 1% FDR of the PEAKS DB result for establishing a confidence threshold for PSMs.

Data processing was performed using Perseus Software (version 2.0.9.0) with default settings as workflow depicted in **Supplemental Fig. S6**. Intensity values were log2-transformed, and the quantitative profiles were filtered for missing values as identified proteins had to be present in at least 70% of samples for further analysis. Data normalization was performed by dividing each protein intensity by the total intensity sum of all identified proteins in its sample, respectively. Significantly different protein groups were assessed by pairwise comparison analyses with Student’s T-test with 0.05 probability level with Benjamini-Hochberg FDR correction. Hierarchical clustering was based on Euclidean distance and on the average linkage of 300 clusters for both row and column trees. The mass spectrometry proteomics data have been deposited to the ProteomeXchange Consortium via the PRIDE (Perez-Riverol et al., 2022) partner repository with the dataset identifier PXD047255 (pellet) and PXD047263 (upper phase).

### RNA extraction and quantitative real-time PCR

Five milliliter of cell cultures were harvested by centrifugation (1 min, 20°C, 1,790 *g*) and flash frozen in liquid nitrogen and conserved at -80°C. Total RNA was extracted using the Direct-Zol™ RNA MiniPrep (Zymo Research, Irvine, CA, USA) and treated with the RNase-free DNase Set (Zymo Research) according to the manufacturer’s instructions.

In order to quantitative real-time PCR (qRT-PCR), complementary DNA (cDNA) was synthesized from 1 µg of total RNA using SuperScript VILO Master Mix (Invitrogen). qRT-PCR was performed on a 480 LightCycler thermocycler (Roche, Indianapolis, IN, USA) following the manufacturer’s instructions with TB Green Premix Ex Taq II (Takara, Shiga, Japan) and the primers listed in **Supplemental Table S7**.

### RNA sequencing

The RNA-seq analysis was carried out on 4 replicates for the 137AH strain and the *dyrkp* mutant. A cDNA library was constructed from 1 µg of total RNA and Illumina HiSeq 2500 sequencing was performed at the Biopuces et Sequencage platform (Illkirch, France). In general, between 28 to 37 million 50-nt single-end reads were generated for each replicate (**Supplemental Table S5**). We processed the fastq files with trimgalore in search of adapters and to trim low-quality nucleotides and remove reads shorter than 20 bp after removing adapters and low-quality bases. Then, we used ribodetector to detect and remove any rRNA that might have survived polyA selection/ribodepletion. The remaining reads were aligned onto the *C. reinhardtii* genome assembly version v6.0 using the Bowtie2 software. HTSeq count was used to identify uniquely mapped reads. For each replicate, between 19 and 33 million reads were mapped on the *C. reinhardtii* genome. DESeq2 package was used to analyze differential gene expression. Only genes with a *p*-value adjusted ≤ 0.01 and an absolute log2 fold change (log2FC) ≥ 1 or ≤ -1 were kept for further analysis. Of 16,288 identified transcripts, 2,139 were categorized as Differentially Expressed Genes (DEGs) (**Supplemental Table S5**). The raw data of RNA-Seq was submitted to NCBI under the accession number (BioProject: PRJNA1042620). The gene ontology term annotation for molecular function and biological process as well as enrichment analysis of DEGs was conducted using the functional enrichment analysis of the STRING database (version 10.0) with default parameters (Szklarczyk et al., 2014). The gene annotation was cross-checked by searching manually on the Phytozome server (*C. reinhardtii* v5.6).

## Acknowledgements

This work was supported by the French ANR grant (TOR-DYRKcontrol) (Y.L.-B.). The use of the HelioBiotec and Zoom platforms of the BIAM institute is also acknowledged. We thank Pascaline Auroy for preparation of samples for RNA sequencing, and Ignasi Andreu Godall from the Bioinformatics Core Unit at CRAG for RNA-seq mapping to *C. reinhardtii* genome assembly.

## Authors contributions

M.K. and Y.L.-B. conceived the study. M.K. and Y.L.-B. designed the experiments and interpreted the results. M.K., S.C., M.B., G.L.J., J.J.T., M.M., M.A., and J.-S.Y. performed experiments. M.K., S.C., M.A., and M.B. visualized the data. Y.L.-B. took part in funding acquisition and administrated the study. F.B., G.P., and Y.L.-B. supervised and commented on the study. M.K. and Y.L.-B. wrote the manuscript with comments from other authors.

## Data and material availability

All biological material described in this study is available upon request. All data are available in the main text or the supplementary materials.

## References

1. Anderson, C.T., and Kieber, J.J. (2020). Dynamic construction, perception, and remodeling of plant cell walls. Annual Review of Plant Biology 71, 39–69.

2. Aranda, S., Laguna, A., and Luna, S.d.l. (2011). DYRK family of protein kinases: evolutionary relationships, biochemical properties, and functional roles. The FASEB Journal 25, 449–462.

3. Bashline, L., Lei, L., Li, S., and Gu, Y. (2014). Cell wall, cytoskeleton, and cell expansion in higher plants. Molecular Plant 7, 586–600.

4. Baudelet, P.-H., Ricochon, G., Linder, M., and Muniglia, L. (2017). A new insight into cell walls of Chlorophyta. Algal Research 25, 333–371.

5. Becker, W. (2012). Emerging role of DYRK family protein kinases as regulators of protein stability in cell cycle control. Cell Cycle 11, 3389–3394.

6. Bonnans, C., Chou, J., and Werb, Z. (2014). Remodelling the extracellular matrix in development and disease. Nature reviews Molecular cell biology 15, 786–801.

7. Brown, J.M., Cochran, D.A., Craige, B., Kubo, T., and Witman, G.B. (2015). Assembly of IFT trains at the ciliary base depends on IFT74. Current Biology 25, 1583–1593.

8. Carmena, M., and Earnshaw, W.C. (2003). The cellular geography of aurora kinases. Nature reviews Molecular cell biology 4, 842–854.

9. Cosgrove, D.J. (2005). Growth of the plant cell wall. Nature reviews molecular cell biology 6, 850–861.

10. Cross, F.R., and Umen, J.G. (2015). The Chlamydomonas cell cycle. The Plant Journal 82, 370–392.

11. de Carpentier, F., Lemaire, S.D., and Danon, A. (2019). When unity is strength: the strategies used by Chlamydomonas to survive environmental stresses. Cells 8, 1307.

12. de Carpentier, F., Maes, A., Marchand, C.H., Chung, C., Durand, C., Crozet, P., Lemaire, S.D., and Danon, A. (2022). How abiotic stress-induced socialization leads to the formation of massive aggregates in Chlamydomonas. Plant Physiology.

13. Domozych, D.S., and LoRicco, J.G. (2023). The extracellular matrix of green algae. Plant Physiology, kiad384.

14. Flinn, B.S. (2008). Plant extracellular matrix metalloproteinases. Functional Plant Biology 35, 1183–1193.

15. Furuya, T., Shinkawa, H., Kajikawa, M., Nishihama, R., Kohchi, T., Fukuzawa, H., and Tsukaya, H. (2021). A plant-specific DYRK kinase DYRKP coordinates cell morphology in Marchantia polymorpha. Journal of plant research 134, 1265–1277.

16. Golldack, D., Popova, O.V., and Dietz, K.-J. (2002). Mutation of the matrix metalloproteinase At2-MMP inhibits growth and causes late flowering and early senescence in Arabidopsis. Journal of Biological Chemistry 277, 5541–5547.

17. Goodenough, U., and Lee, J.-H. (2023). Cell walls. In The Chlamydomonas Sourcebook (Elsevier), pp. 41–64.

18. Gu, Y., and Rasmussen, C.G. (2022). Cell biology of primary cell wall synthesis in plants. The Plant Cell 34, 103–128.

19. Holmbeck, K., Bianco, P., Caterina, J., Yamada, S., Kromer, M., Kuznetsov, S.A., Mankani, M., Robey, P.G., Poole, A.R., and Pidoux, I. (1999). MT1-MMP-deficient mice develop dwarfism, osteopenia, arthritis, and connective tissue disease due to inadequate collagen turnover. Cell 99, 81–92.

20. Huang, Y., and Thelen, J.J. (2012). KiC assay: a quantitative mass spectrometry-based approach. Quantitative Methods in Proteomics, 359-370.

21. Khona, D.K., Shirolikar, S.M., Gawde, K.K., Hom, E., Deodhar, M.A., and D’Souza, J.S. (2016). Characterization of salt stress-induced palmelloids in the green alga, Chlamydomonas reinhardtii. Algal Research 16, 434–448.

22. Kim, J., Lee, S., Baek, K., and Jin, E. (2020). Site-specific gene knock-out and on-site heterologous gene overexpression in Chlamydomonas reinhardtii via a CRISPR-Cas9-mediated knock-in method. Frontiers in plant science 11, 306.

23. Kubo, T., Kaida, S., Abe, J., Saito, T., Fukuzawa, H., and Matsuda, Y. (2009). The Chlamydomonas hatching enzyme, sporangin, is expressed in specific phases of the cell cycle and is localized to the flagella of daughter cells within the sporangial cell wall. Plant and Cell Physiology 50, 572–583.

24. Kumar, B., Koul, S., Petersen, J., Khandrika, L., Hwa, J.S., Meacham, R.B., Wilson, S., and Koul, H.K. (2010). p38 mitogen-activated protein kinase–driven MAPKAPK2 regulates invasion of bladder cancer by modulation of MMP-2 and MMP-9 activity. Cancer research 70, 832–841.

25. Kurabayashi, N., and Sanada, K. (2013). Increased dosage of DYRK1A and DSCR1 delays neuronal differentiation in neocortical progenitor cells. Genes & development 27, 2708–2721.

26. Laguna, A., Aranda, S., Barallobre, M.J., Barhoum, R., Fernández, E., Fotaki, V., Delabar, J.M., de la Luna, S., de la Villa, P., and Arbonés, M.L. (2008). The protein kinase DYRK1A regulates caspase-9-mediated apoptosis during retina development. Developmental cell 15, 841–853.

27. Li, X., Patena, W., Fauser, F., Jinkerson, R.E., Saroussi, S., Meyer, M.T., Ivanova, N., Robertson, J.M., Yue, R., and Zhang, R. (2019). A genome-wide algal mutant library and functional screen identifies genes required for eukaryotic photosynthesis. Nature genetics 51, 627–635.

28. Long, H., Zhang, F., Xu, N., Liu, G., Diener, D.R., Rosenbaum, J.L., and Huang, K. (2016). Comparative analysis of ciliary membranes and ectosomes. Current Biology 26, 3327–3335.

29. Mishra, L.S., Kim, S.Y., Caddell, D.F., Coleman-Derr, D., and Funk, C. (2021). Loss of Arabidopsis matrix metalloproteinase-5 affects root development and root bacterial communities during drought stress. Physiologia Plantarum 172, 1045–1058.

30. Noh, E.-M., Lee, Y.-R., Hong, O.-Y., Jung, S.H., Youn, H.J., and Kim, J.-S. (2015). Aurora kinases are essential for PKC-induced invasion and matrix metalloproteinase-9 expression in MCF-7 breast cancer cells. Oncology reports 34, 803–810.

31. Page-McCaw, A., Ewald, A.J., and Werb, Z. (2007). Matrix metalloproteinases and the regulation of tissue remodelling. Nature reviews Molecular cell biology 8, 221–233.

32. Perez-Riverol, Y., Bai, J., Bandla, C., García-Seisdedos, D., Hewapathirana, S., Kamatchinathan, S., Kundu, D.J., Prakash, A., Frericks-Zipper, A., and Eisenacher, M. (2022). The PRIDE database resources in 2022: a hub for mass spectrometry-based proteomics evidences. Nucleic acids research 50, D543–D552.

33. Proost, S., and Mutwil, M. (2018). CoNekT: an open-source framework for comparative genomic and transcriptomic network analyses. Nucleic acids research 46, W133–W140.

34. Schulz-Raffelt, M., Chochois, V., Auroy, P., Cuiné, S., Billon, E., Dauvillée, D., Li-Beisson, Y., and Peltier, G. (2016). Hyper-accumulation of starch and oil in a Chlamydomonas mutant affected in a plant-specific DYRK kinase. Biotechnology for biofuels 9, 1–12.

35. Seifert, G.J., and Blaukopf, C. (2010). Irritable walls: the plant extracellular matrix and signaling. Plant physiology 153, 467–478.

36. Shoshan, E., Braeuer, R.R., Kamiya, T., Mobley, A.K., Huang, L., Vasquez, M.E., Velazquez-Torres, G., Chakravarti, N., Ivan, C., Prieto, V., Villares, G.J., and Bar-Eli, M. (2016). NFAT1 Directly Regulates IL8 and MMP3 to Promote Melanoma Tumor Growth and Metastasis. Cancer Research 76, 3145–3155.

37. Shou, J., Jing, J., Xie, J., You, L., Jing, Z., Yao, J., Han, W., and Pan, H. (2015). Nuclear factor of activated T cells in cancer development and treatment. Cancer letters 361, 174–184.

38. Szklarczyk, D., Franceschini, A., Wyder, S., Forslund, K., Heller, D., Huerta-Cepas, J., Simonovic, M., Roth, A., Santos, A., Tsafou, K.P., Kuhn, M., Bork, P., Jensen, L.J., and von Mering, C. (2014). STRING v10: protein–protein interaction networks, integrated over the tree of life. Nucleic Acids Research 43, D447–D452.

39. Tryggvason, K., Höyhtyä, M., and Salo, T. (1987). Proteolytic degradation of extracellular matrix in tumor invasion. Biochimica et Biophysica Acta (BBA)-Reviews on Cancer 907, 191–217.

40. Wagner, E.F., and Nebreda, A.R. (2009). Signal integration by JNK and p38 MAPK pathways in cancer development. Nature Reviews Cancer 9, 537–549.

41. Wang, L., Wen, X., Wang, Z., Lin, Z., Li, C., Zhou, H., Yu, H., Li, Y., Cheng, Y., Chen, Y., Lou, G., Pan, J., and Cao, M. (2022). Ciliary transition zone proteins coordinate ciliary protein composition and ectosome shedding. Nature Communications 13, 3997.

42. Wilkinson, D.J., Desilets, A., Lin, H., Charlton, S., del Carmen Arques, M., Falconer, A., Bullock, C., Hsu, Y.-C., Birchall, K., Hawkins, A., Thompson, P., Ferrell, W.R., Lockhart, J., Plevin, R., Zhang, Y., Blain, E., Lin, S.-W., Leduc, R., Milner, J.M., and Rowan, A.D. (2017). The serine proteinase hepsin is an activator of pro-matrix metalloproteinases: molecular mechanisms and implications for extracellular matrix turnover. Scientific Reports 7, 16693.

43. Wood, C.R., Huang, K., Diener, D.R., and Rosenbaum, J.L. (2013). The cilium secretes bioactive ectosomes. Current Biology 23, 906–911.

44. Wu, H., Si, Q., Liu, J., Yang, L., Zhang, S., and Xu, J. (2022). Regulation of Arabidopsis Matrix Metalloproteinases by Mitogen-Activated Protein Kinases and Their Function in Leaf Senescence. Frontiers in plant science 13.

45. Zhang, J., Xin, L., Shan, B., Chen, W., Xie, M., Yuen, D., Zhang, W., Zhang, Z., Lajoie, G.A., and Ma, B. (2012). PEAKS DB: de novo sequencing assisted database search for sensitive and accurate peptide identification. Molecular & cellular proteomics 11.

46. Zou, Y., and Bozhkov, P.V. (2021). Chlamydomonas proteases: classification, phylogeny, and molecular mechanisms. Journal of Experimental Botany 72, 7680–7693.

